# Consumption of a Western-style diet modulates the response of the murine gut microbiome to ciprofloxacin

**DOI:** 10.1101/780049

**Authors:** Damien J. Cabral, Jenna I. Wurster, Benjamin J. Korry, Swathi Penumutchu, Peter Belenky

## Abstract

Dietary composition and antibiotic use are known to have major impacts on the structure and function of the gut microbiome, often resulting in dysbiosis. Despite this, little research has been done to explore the role of host diet as a determinant of antibiotic-induced microbiome disruption.

Here, we utilize a multi-omic approach to characterize the impact of Western-style diet consumption on ciprofloxacin-induced changes to gut microbiome community structure and transcriptional activity. We found that mice consuming a Western-style diet experienced a greater expansion of *Firmicutes* following ciprofloxacin treatment than those eating a control diet. At the transcriptional level, we found that ciprofloxacin induced a reduction in the abundance of TCA cycle transcripts on both diets, suggesting that carbon metabolism plays a key role in the response of the gut microbiome to this antibiotic. Despite this shared response, we observed extensive differences in the response of the microbiota to ciprofloxacin on each diet. In particular, at the whole-community level we detected an increase in starch degradation, glycolysis, and pyruvate fermentation following antibiotic treatment in mice on the Western diet, which we did not observe in mice on the control diet. Similarly, we observed diet-specific changes in the transcriptional activity of two important commensal bacteria, *Akkermansia muciniphila* and *Bacteroides thetaiotaomicron*, involving diverse cellular processes such as nutrient acquisition, stress responses, and capsular polysaccharide (CPS) biosynthesis. These findings demonstrate that host diet plays a key role in determining the extent of disruption of microbiome composition and function induced by antibiotic treatment.

**Importance:** While both diet and antibiotics are individually known to have profound impacts on gut microbiome composition, little work has been done to examine the effect of these two factors combined. A number of negative health outcomes, including diabetes and obesity, are associated with diets high in simple sugars in fats but low in host-indigestible fiber, and some of these outcomes may be mediated by the gut microbiome. Likewise, treatment with broad-spectrum antibiotics and the resulting dysbiosis is associated with many of the same detrimental side effects. Previous work has shown that nutrient availability, as influenced by host diet, plays an important role in determining the extent of antibiotic-induced disruption to the gut microbiome. Due to the growing incidence of disorders related to antibiotic-induced dysbiosis, it is essential to determine how the prevalence of high fat and sugar “Western”-style diets impacts the response of the microbiome to antibiotics.

## Background

The gut microbiome includes the trillions of largely commensal bacteria, archaea, fungi, and viruses and their collective genetic material that inhabit the gastrointestinal tract (1-3). These communities play an important role in numerous biological processes, such as digestion, neurological development, colonization resistance, and immune function (4-18). However, the gut microbiome is exquisitely sensitive to perturbation and the disruption of microbial homeostasis, known as dysbiosis, can result in numerous harmful impacts to the host. In particular, broad-spectrum antibiotic use is known to have numerous detrimental impacts on the gut microbiota. Within hours of treatment, antibiotics induce dramatic reductions in both bacterial loads and diversity within the microbiome (19, 20).

While compositional changes are typically transient and recover following the cessation of therapy, oftentimes the structure and diversity of the microbiota never return to their pre-treatment levels. These changes often result in dysbiosis and have numerous acute and chronic impacts on host health. In particular, dysbiosis may increase the risk of infection with opportunistic fungal and bacterial pathogens by reducing colonization resistance (1, 5-7, 21-25). Most notably, broad-spectrum antibiotic treatment is known to be a major risk factor in *Clostridioides difficile* infection, which is responsible for approximately 29,000 deaths worldwide and 15,000 in the United States alone (21, 23, 26, 27). In addition to short-term complications such as pathogen blooms, persistent dysbiosis is correlated with a number of chronic conditions associated with considerable morbidity and mortality, such as asthma, obesity, and inflammatory bowel disease (5, 8-10, 13, 15, 17, 18, 28).

In addition to eliciting changes in community structure, antibiotic exposure also has a dramatic impact on the functional activity of the gut microbiome by altering the expression of key metabolic genes and the availability of carbohydrates within the gut (20). Most notably, amoxicillin treatment has been shown to increase the expression of glycoside hydrolases responsible for hydrolysis of polysaccharides, while simultaneously decreasing the abundance of transcripts encoding sugar phosphotransferase systems (PTS) responsible for uptake of simple sugars (20). Reflecting these changes, amoxicillin also decreases the concentration of glucose within the ceca of mice (20). Furthermore, dietary intervention experiments have demonstrated that the response of the microbiota to antibiotics can be impacted by nutrient modulation. For example, glucose supplementation reduces the absolute abundance of bacteria, particularly *Bacteroides thetaiotaomicron*, following amoxicillin treatment in mice (20). Therefore, it is likely that host diet composition has a major impact on the response of these communities to perturbation.

It is also known that dietary composition has a profound impact on microbiome diversity and overall gut health (29-35). Diets high in fat and simple sugars, typically referred to as “Western” diets, have been associated with a number of negative health states including obesity, diabetes, allergies, and inflammatory bowel disease (36-46). Such diets have very low levels of microbiota-accessible carbohydrates (MACs), which are typically found in complex plant polysaccharides and are indigestible and unabsorbable by the host (40, 44, 47-49). MACs are typically fermented by the colonic microbiota to produce short-chain fatty acids (SCFAs), which in turn play important roles in regulating energy homeostasis and inflammation within the host (40, 45, 50-55). In addition to being associated with increased levels of SCFAs, high-MAC diets have been shown to increase microbial diversity, a classic benchmark for gut microbiota health. Conversely, low-MAC Western diets are known to reduce both microbiome diversity and SCFA production (44, 46, 49, 56). Due to the absence of MACs, such diets also enrich for muciniphilic microbes that may degrade mucosal lining of the gut, such as *Akkermansia muciniphila* (40, 42, 48). Degradation of the mucosal layer over time is hypothesized to result in compromised gut barrier function that may ultimately lead to increased inflammation and colitis. Lastly, MAC-deficient diets have been shown to exacerbate infection with *C. difficile*, suggesting that they may negatively impact colonization resistance (26, 27).

Individually, antibiotic usage and the consumption of Western-style diets are known to negatively impact the microbiota, resulting in similar long-term impacts on host health (including obesity, allergies, and inflammatory bowel disease). Despite this, little work has been done to explore how diet impacts the response of the microbiota to antibiotics. Previous work has suggested that dietary composition may play an important role in determining the extent of antibiotic-induced microbiome disruption (20). In this study, we use a combined metagenomic and metatranscriptomic approach to characterize the impact of a Western-style diet on the taxonomic and functional disruption of the microbiome during ciprofloxacin treatment. Using shotgun metagenomics, we found that ciprofloxacin elicited differential impacts on community composition in mice at both the phylum- and species-level that were diet-dependent. Using metatranscriptomics, we observed that consumption of a Western diet itself induced profound transcriptional changes within the gut microbiomes of mice; furthermore, consumption of this diet modulated the transcriptional response of these communities to antibiotic treatment. In particular, dietary composition had a major impact on the abundance of transcripts encoding key metabolic genes. Lastly, we were able to detect unique species-specific transcriptional changes in response to both diet and ciprofloxacin treatment in two important commensal bacteria, *A. muciniphila* and *B. thetaiotaomicron*.

## Results

To determine the impact of dietary composition and antibiotic exposure on the structure and function of the murine gut microbiome, female C57BL/6J mice were randomly assigned to either a high-fat, high-sugar “Western”-style diet, or a low-fat control diet for seven days in multiple cages. At this point, mice from each diet were again randomly split between ciprofloxacin and vehicle control groups and treated for 24 hours in multiple cages (n = 8-to-12 per group). This time point was chosen because previously published work demonstrated that 24 hours of ciprofloxacin treatment was sufficient to induce changes in community structure and transcriptional activity (20). Additionally, this time frame allowed us to profile the acute response of the microbiota to ciprofloxacin exposure, rather than characterizing a post-antibiotic state of equilibrium. Following treatment, the mice were sacrificed to harvest their cecal contents for taxonomic profiling and transcriptional analysis (Figure 1A). Overall, we found that diet and ciprofloxacin treatment had a significant impact on gut microbiome structure (Figure 1B+C, Figure S1).

**Figure 1:**
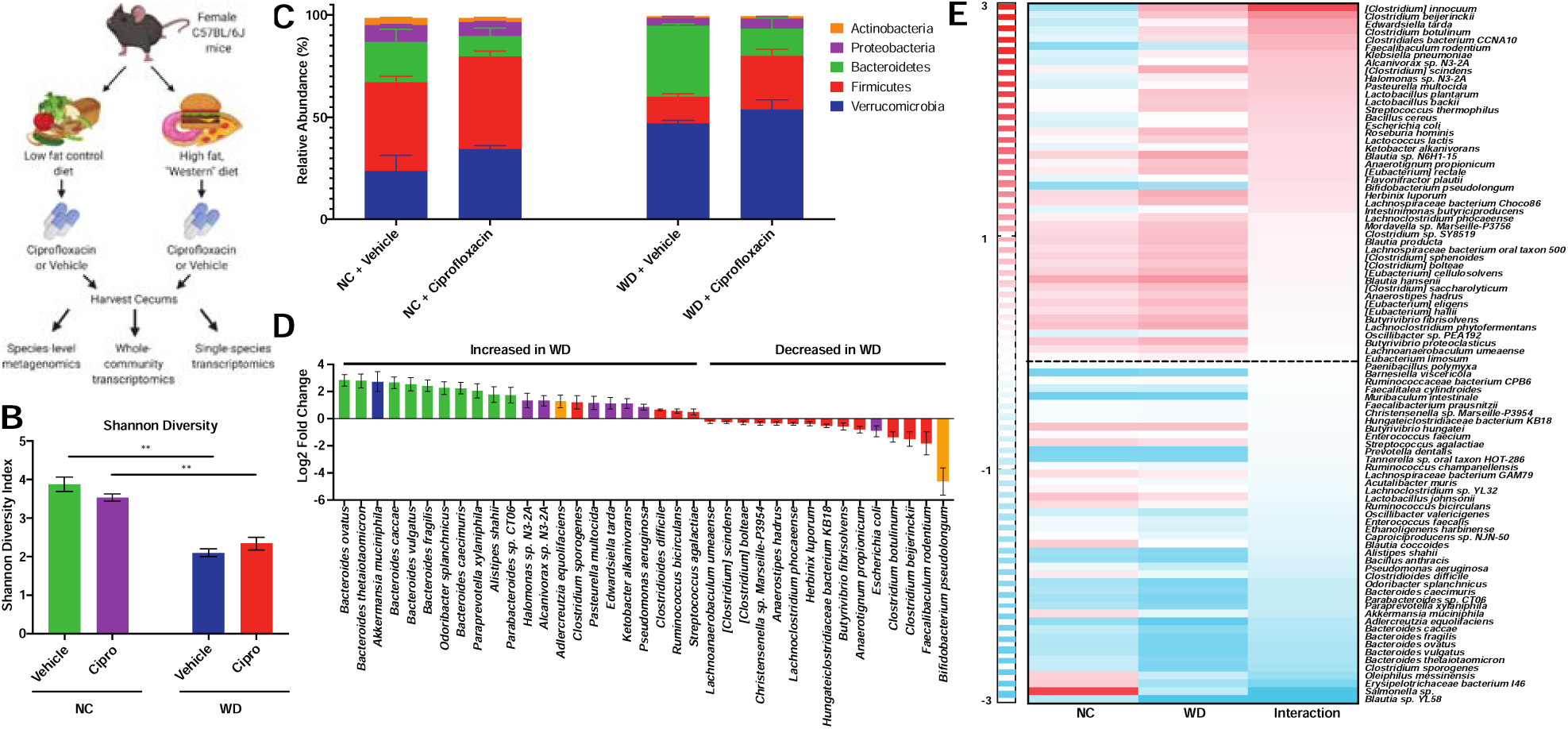
Impact of diet and ciprofloxacin administration on murine gut microbiome composition. A. Experimental workflow used in this study. Figure was created with Biorender.com. B. Alpha diversity of experimental groups as measured by the Shannon Diversity Index. Data are represented as mean ± standard error of the mean (SEM). (**p<0.01, Welch ANOVA with Dunnett T3 test for multiple hypothesis testing). C. Stacked barplot of the five most abundant bacterial phyla in our dataset. Data are represented as mean + SEM for each phylum. D. Differentially abundant (Benjamini-Hochberg adjusted p-value < 0.1) bacterial species detected in mice consuming the Western diet (WD). Data are represented as log_2_ fold change relative to control diet ± standard error. Bar color denotes phylum level taxonomic classification (blue – *Verrucomicrobia*, red – *Firmicutes*, green – *Bacteroidetes*, purple – *Proteobacteria*, orange – *Actinobacteria*). E. Heatmap of the change in abundance of the top 90 bacterial species in response to ciprofloxacin on control (NC) and Western (WD) diets. The Interaction column represents the interaction term generated by DESeq2, denoting the impact of diet on the change in abundance of each species to ciprofloxacin. Cell color denotes log_2_ fold change of a particular species in response to ciprofloxacin. Heatmap rows were sorted by interaction term value from highest to lowest. For all analyses, n = 4. For full DESeq2 results, see Additional File 1.

We first assessed the effects that diet and ciprofloxacin have on the diversity of the gut microbiome using 16S ribosomal RNA (rRNA) sequencing. Mice consuming the Western diet had significantly less diverse gut microbiomes than those fed the control diet (Figure S1A). Interestingly, we also observed that the Western diet exacerbated the reduction in alpha diversity during ciprofloxacin treatment (Figure S1A). Next, we performed principle coordinate analysis using Bray-Curtis dissimilarity paired with PERMANOVA analysis to profile the degree of dissimilarity between our samples and the significance of this distance. Unsurprisingly, our samples formed four distinct clusters, driven by both diet and ciprofloxacin treatment (Figure S1B).

Due to the limited phylogenetic resolution provided by 16S rRNA sequencing, in addition to its inability to provide functional information about sequenced communities, we opted to perform shotgun metagenomics and metatranscriptomic analysis on a subset of our samples, representing mice from multiple cages (n = 4 per treatment group) (20, 57-60). Using metagenomics, we observed that consumption of the Western diet reduced community diversity (Figure 1B). Taxonomically, mice fed a Western diet displayed elevated levels of the phyla *Verrucomicrobia* and *Bacteroidetes*, and a reduced abundance of *Firmicutes* (Figure 1C). At the species level, these shifts appear to be largely driven by an expansion of members of the *Bacteroides* genus (Figure 1D, Figure S1). Additionally, the Western diet-fed mice displayed an elevated abundance of several species from the *Proteobacteria* phylum, which has been previously shown to be suggestive of dysbiosis (61). Two important bacterial species found in the gut microbiomes of both mice and humans, *B. thetaiotaomicron* and *A. muciniphila*, were observed at significantly elevated levels in the mice fed a Western diet (Figure 1D, Figure S1). Notably, both species are known to utilize host-produced mucins; thus, this observation is consistent with earlier studies that have suggested that the consumption of a low-MAC Western diet enriches for muciniphilic bacteria (40, 42, 48).

Overall, host diet appears to have a major impact on the structure of the microbiome during ciprofloxacin treatment. Although ciprofloxacin did not induce a significant reduction in alpha diversity in the timeframe tested, we found that diet drove differential community composition following antibiotic exposure. At the phylum level we observed a significant expansion in the relative abundance of *Firmicutes* following ciprofloxacin treatment on the Western diet (adjusted p-value = 0.0388) but not on the control diet (adjusted p-value = 0.8718) (Figure 1C). To determine which species displayed a differential response to ciprofloxacin on the Western and control diets, we utilized DESeq2 to analyze the interaction between diet and antibiotic treatment (62). In our analysis, an interaction term describes the direction and magnitude of the differential impact that host diet has on antibiotic perturbation within the microbiome. For example, ciprofloxacin reduces the abundance of *Clostridium innocuum* on the control diet, while it expands following treatment on the Western diet (Figure 1E, Additional File 1); thus, the interaction term in this case is positive, as it indicates an expansion on the Western diet relative to control following ciprofloxacin exposure.

While most species responded similarly to ciprofloxacin therapy on both diets, there were several notable exceptions. For example, the expansion of several *Clostridium* species (such as *C. innocuum, Clostridium beijerinckii*, and *Clostridium scindens*) following ciprofloxacin tended to be higher on the Western diet than the control (denoted by a positive interaction value in Figure 1E, Additional File 1). Conversely, the reduction of several *Bacteroides* species following antibiotic treatment tended to be exacerbated on the Western diet (negative interaction values, Figure 1E, Additional File 1). In particular, the reduction of *B. thetaiotaomicron* in response to ciprofloxacin was enhanced on the Western diet; in contrast, dietary composition had no significant impact on the response of *A. muciniphila* to ciprofloxacin (Additional File 1).

### Ciprofloxacin Elicits Unique Shifts in Gene Expression on Western and Control Diets

We detected clear differences in the susceptibility of the microbiome to treatment on the two diets. It is possible that the differential metabolic environment between the two diets changes the activity of the gut microbiome and contributes to altered susceptibility. Though many studies have examined the impacts of either diet or antibiotic treatment on the gut microbiome, few have examined their combined effect on the composition or gene expression of these communities. To address this, we first analyzed our metatranscriptomic dataset using the HUMAnN2 pipeline, which normalizes the abundance of RNA transcripts against their corresponding gene abundance in the metagenomic data (63). Thus, this tool normalizes for differences in community composition between experimental groups and facilitates the comparison of metabolic pathway expression at the whole-community level. A comparison of the transcriptional profile of all experimental groups demonstrates that the microbiota of mice consuming the Western diet display elevated expression of tricarboxylic acid (TCA) cycle and fatty acid degradation pathways in both vehicle and ciprofloxacin treatments, likely reflective of the increased fat and sugar content of this diet (Figure 2A, Additional File 2). Additionally, we found an elevated expression of glycogen degradation genes in the Western diet mice receiving ciprofloxacin, which was not observed in any other group (Figure 2A, Additional File 2). Conversely, the microbiota of mice consuming the control diet appeared to have elevated expression of amino acid biosynthesis pathways (namely isoleucine, aspartate, asparagine, lysine, and histidine) regardless of antibiotic treatment (Figure 2A, Additional File 2). Interestingly, we also observed elevated levels of several different pathways of nucleotide biosynthesis in the control diet samples receiving vehicle while the Western diet mice displayed elevated levels of adenosine and guanosine nucleotide degradation (Figure 2A, Additional File 2).

**Figure 2:**
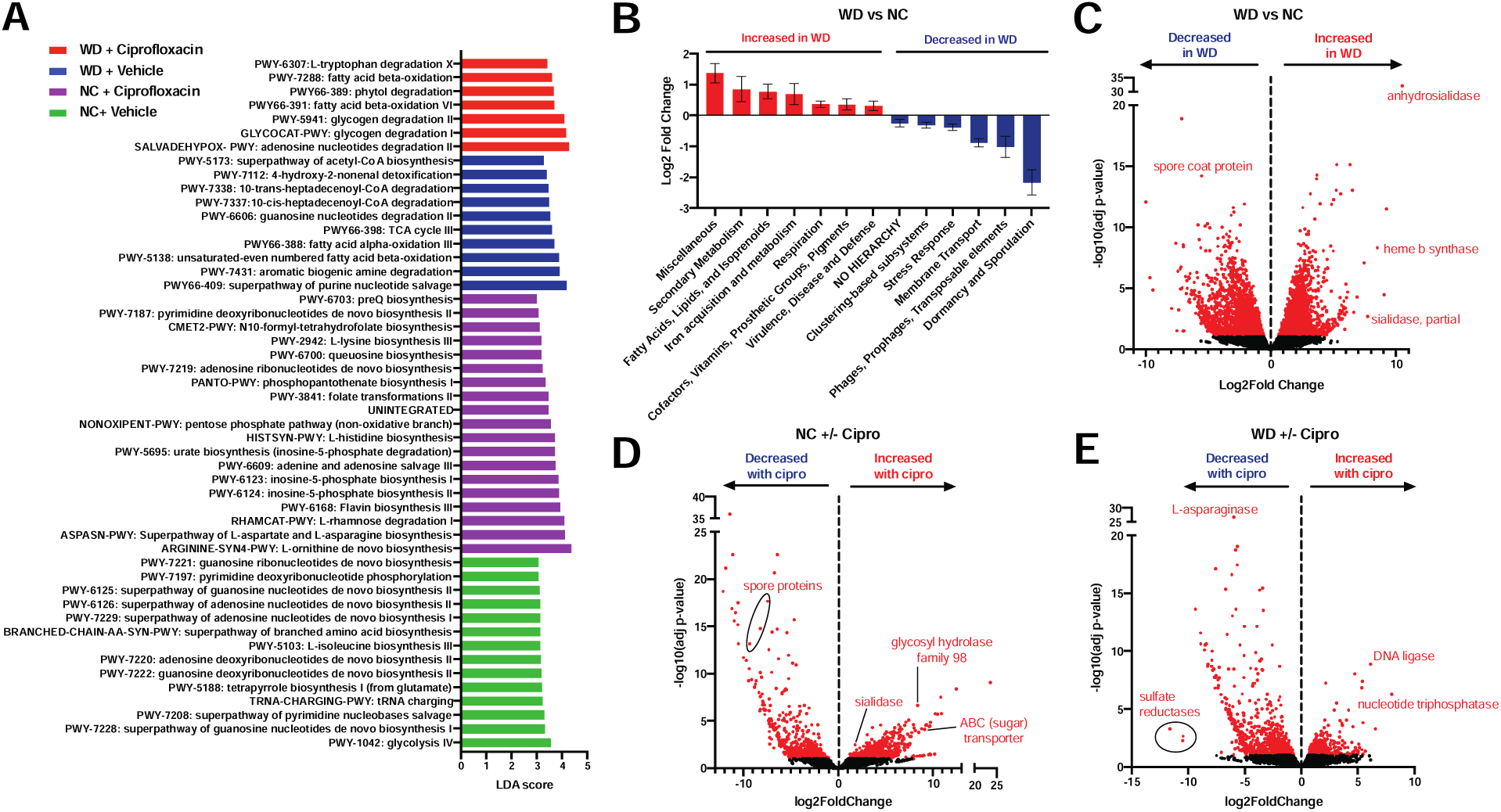
Ciprofloxacin elicits unique shifts gene expression on Western and control diets at the MetaCyc pathway level. A. Linear discriminant analysis (LDA) of MetaCyc pathways that were differentially associated with each experimental group. Bar size indicates LDA score and color indicates the experimental group that a MetaCyc pathway was significantly associated with. All LDA scores were generated using LEfSe on unstratified pathway outputs from HUMAnN2. For full pathway names and statistics, see Additional File 2. B. Differentially expressed (Benjamini-Hochberg adjusted p-value < 0.1) level 1 SEED subsystems in the murine cecal metatranscriptome in ciprofloxacin-treated mice consuming the control (NC) diet. Data are represented as log_2_ fold change relative to vehicle control ± standard error. Only features with a base mean ≥ 50 were plotted. See Additional File 3 for full results. (C – E) Volcano plots of the metatranscriptomic profile of the murine cecal microbiome in vehicle-treated mice consuming Western diet (C), ciprofloxacin-treated mice on the control diet (D), and ciprofloxacin-treated mice on the Western diet (E). Data was generated by aligning metatranscriptomic reads to RefSeq using SAMSA2 and analyzing using DESeq2. Points in red represent transcripts for which a statistically significant change in expression was detected (Benjamini-Hochberg adjusted p-value < 0.1). Select genes of interest are labeled. See Additional File 4 for full results. For all analyses, n = 4.

A pairwise comparison between the vehicle-treated samples on the Western and control diets reveals that the microbiota exhibits extensive transcriptional changes in response to dietary modulation (Additional File 2). Notably, the microbiota on the control diet-fed mice displayed elevated expression of nucleotide biosynthesis, glycolysis, gluconeogenesis, starch degradation, and pyruvate fermentation (Figure 2A, Additional File 2). We also observed increased expression of the *Bifidobacterium* shunt, which is known to play a role in SCFA production and may provide mechanistic insight into the reduced SCFA levels observed on the Western diet in other studies (40, 51) (Additional File 2).

When we compared the response of the microbiome to ciprofloxacin on each diet, we found key differences in the overall transcriptional profiles (Additional File 2). In mice consuming the Western diet, ciprofloxacin treatment was associated with increased abundance of transcripts from glycogen and starch degradation, glycolysis, and pyruvate fermentation (Additional File 2). Notably, the expression of glycogen degradation was elevated in vehicle-treated samples on the control diet, suggesting that the utilization of this pathway during ciprofloxacin treatment is diet-dependent (Additional File 2). On the control diet, we observed an increased abundance of inosine-5-phosphate biosynthesis with ciprofloxacin treatment (Figure 2A, Additional File 2). Interestingly, we observed that TCA cycle expression was reduced in ciprofloxacin-treated mice compared to the vehicle treatment in both control and Western diet conditions – the lone commonality between diets (Additional File 2). Previous work has demonstrated that elevated TCA cycle activity increases sensitivity to bactericidal antibiotics, including fluoroquinolones, *in vitro* (64-68). Thus, this result suggests that TCA cycle activity may play a key role in the response of the microbiota to ciprofloxacin treatment *in vivo*, though more work is required to understand its impact.

### Ciprofloxacin has a differential impact on the abundance of iron metabolism and mucin degradation transcripts on the Western versus control diets

Due to the potential limitations of the use of a single pipeline, we analyzed our metatranscriptomic dataset with SAMSA2 in parallel with HUMAnN2. While HUMAnN2 normalizes for DNA abundance, SAMSA2 does not have this feature and thus the output is representative of overall transcript levels rather than relative expression. Despite these differences, many of the changes observed using HUMAnN2 were detected using SAMSA2 at the SEED subsystem level. When comparing the vehicle-treated samples on both diets, SAMSA2 detected an increased abundance of transcripts related to respiration in the Western diet, mirroring the increase in TCA cycle expression found with HUMAnN2 (Figure 2B, Additional File 3). The microbiota from the Western diet-fed mice also displayed an increased abundance of transcripts involving fatty acids and iron acquisition, which likely reflect altered nutrient availability (Figure 2B, Additional File 3). Furthermore, transcripts related to virulence and disease were elevated in mice consuming the Western diet, further supporting the hypothesis that consumption of a Western diet may promote dysbiosis and the expansion of enteric pathogens (Figure 2B, Additional File 3). Interestingly, comparatively few subsystems were changed in abundance following ciprofloxacin treatment on either diet (Additional File 3). Most notably, we observed a decrease in transcripts related to dormancy and sporulation in response to ciprofloxacin on both diets (Additional File 3). A similar finding was observed in a recent study in which mice were treated with ciprofloxacin while consuming a non-purified diet, suggesting that these transcripts may play a key role in the response of the microbiota to this antibiotic (20).

An additional strength of the SAMSA2 pipeline is that it enables differential abundance testing of individual transcripts in addition to pathway- and subsystem-level analysis (69). We observed that the microbiota of the mice consuming the Western diet displayed increased abundance of heme b synthase transcripts, which may suggest increased iron acquisition (Figure 2C, Additional File 4). Interestingly, in the untreated mice, we detected large, Western diet-associated increases in the abundance of two different transcripts encoding sialidases, which play a key role in the utilization of host-produced mucins (70) (Figure 2C, Additional File 4). While other studies have shown that the consumption of a Western diet enriches for muciniphilic taxa, this observation demonstrates that such a diet also increases transcriptional activity related to mucin degradation within the microbiome (40, 42). Furthermore, ciprofloxacin increased the abundance of sialidase transcripts in mice on the control diet, suggesting that this effect may be exacerbated by antibiotic treatment (Figure 2D, Additional File 4).

In addition to the increased abundance of sialidase transcripts, ciprofloxacin induced a number of notable transcriptional changes on each diet. Reflecting the overall reduction in sporulation seen at the subsystem level, we found that the abundance of several sporulation-related transcripts was reduced on the control diet following ciprofloxacin treatment (Figure 2D, Additional File 4). We also examined the interaction of diet and antibiotic treatment on transcript abundance within the microbiome. Notably, we found that the transcript abundance of several sporulation genes following ciprofloxacin treatment was significantly higher on the Western diet than the control (Additional File 4). Additionally, transcripts encoding phosphotransferase system (PTS) transporters of various substrates (such as cellobiose, fructose, and N-acetylgalactosamine) were also found to be higher on the Western diet following ciprofloxacin treatment (Additional File 4). Conversely, consumption of the Western diet significantly reduced the change in transcript abundance of both pectate lyase and a hemin receptor following ciprofloxacin therapy. Together, these findings demonstrate that dietary composition significantly impacts the transcriptional response of the microbiome to ciprofloxacin.

### Diet and ciprofloxacin alter gene expression within *B. thetaiotaomicron* and *A. muciniphila*

Next we sought to profile how diet and drug treatment impacted the transcriptional response of individual species within the microbiota. We used a previously published pipeline to interrogate the impact of diet and antibiotic treatment on two individual species: *B. thetaiotaomicron* and *A. muciniphila* (20, 71). We focused on these bacteria because they are known human gut commensals, were found in relatively high levels in all samples analyzed, and because they were differentially abundant in a diet-dependent manner.

The Western diet significantly elevated the baseline relative abundance of *A. muciniphila* (Figure 3A). Interestingly, on this diet *A. muciniphila* displayed increased expression of several known stress response genes (Figure 3B, Additional File 5). Specifically, we observed elevated levels of transcripts for catalase HPII (AMUC_RS11055), ATP-dependent chaperone ClpB (AMUC_RS04500), a universal stress protein (AMUC_RS00415), superoxide dismutase (AMUC_RS08510), a UvrB/UvrC protein (AMUC_RS00335), and a thioredoxin family protein (AMUC_RS11695). Together, these changes may indicate that Western diet consumption induces a general stress response in *A. muciniphila*. Additionally, we observed numerous changes within respiratory and central carbon metabolism, suggesting broad metabolic changes in response to the Western diet (Figure 3B, Additional File 5). We specifically detected increased expression of genes encoding terminal oxidases of the respiratory chain (AMUC_RS09050 - cytochrome ubiquinol oxidase subunit I and AMUC_RS09045 - cytochrome d ubiquinol oxidase subunit II), the TCA cycle (AMUC_RS09040 - 2-oxoglutarate dehydrogenase E1 component), glycolysis (AMUC_RS06320 - phosphopyruvate hydratase, AMUC_RS02385 - pyruvate kinase), and pyruvate metabolism (AMUC_RS01195 - ubiquinone-dependent pyruvate dehydrogenase). Of the genes that were significantly reduced on the Western diet, the most notable were an adenylsuccinate synthase (AMUC_RS11340) and a lyase (AMUC_RS10360), which both play important roles in purine metabolism (Figure 3B, Additional File 5) (72).

**Figure 3:**
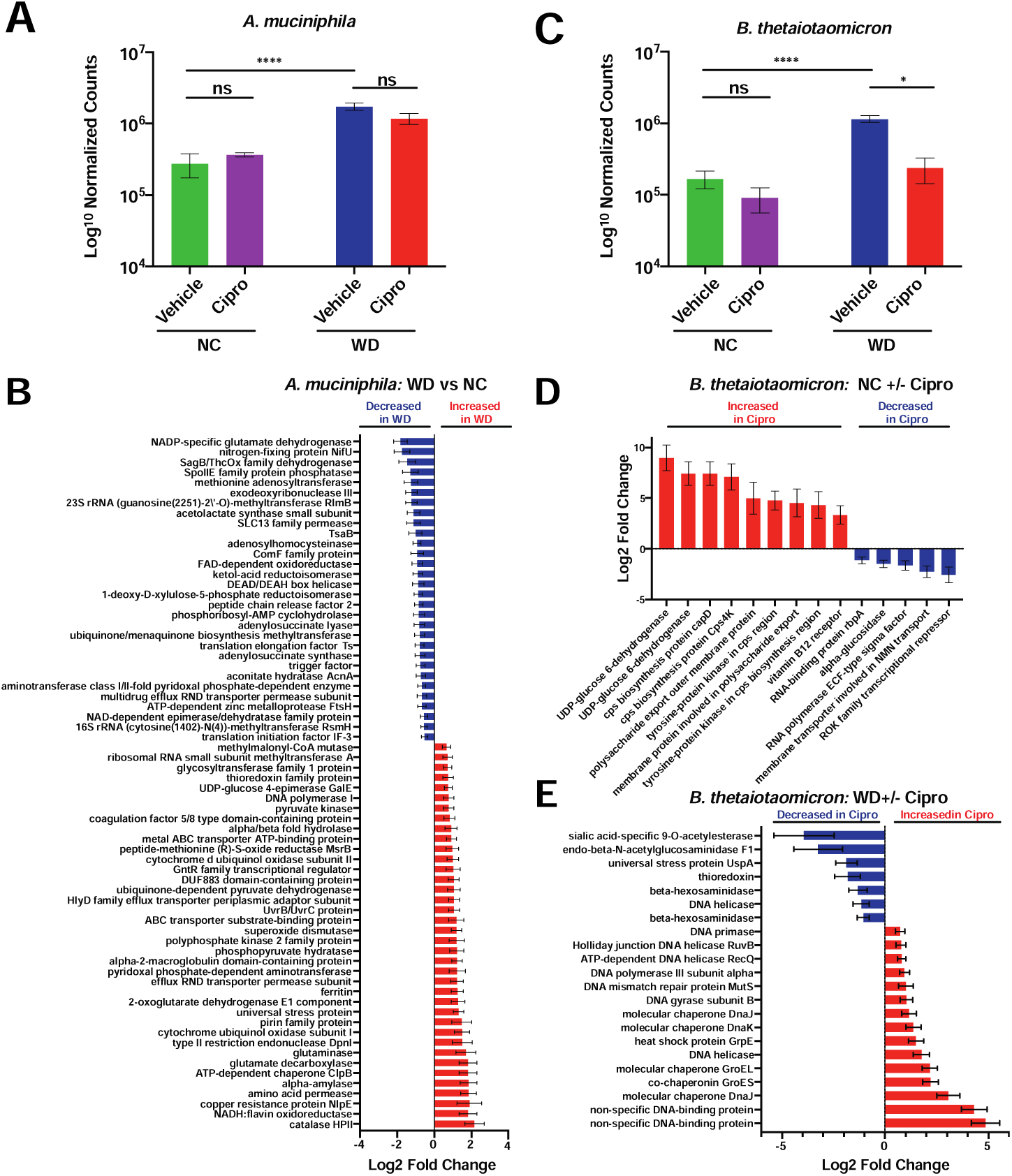
Diet and ciprofloxacin alter gene expression within *B. thetaiotaomicron* and *A. muciniphila*. A. Normalized counts of *A. muciniphila* in each experimental group. Data are represented as mean ± SEM. Normalized counts were generated with DESeq2 and subsequently used to perform differential abundance testing. (*p<0.05, ****p<0.0001, Wald test with Benjamini and Hochberg correction). See Additional File 1 for full results. B. Select differentially expressed (Benjamini-Hochberg adjusted p-value < 0.1) genes of interest in *A. muciniphila* within the cecum of vehicle-treated mice consuming the Western (WD) diet. Data are represented as log_2_ fold change relative to control diet ± standard error. See Additional File 5 for full results. C. Normalized counts of *B. thetaiotaomicron* in each experimental group. Data are represented as mean ± SEM. Normalized counts were generated with DESeq2 and subsequently used to perform differential abundance testing. (*p<0.05, ****p<0.0001, Wald test with Benjamini and Hochberg correction). See Additional File 1 for full results. D. Select differentially expressed (Benjamini-Hochberg adjusted p-value < 0.1) genes of interest in *B. thetaiotaomicron* within the cecum of ciprofloxacin-treated mice consuming the control (NC) diet. Data are represented as log_2_ fold change relative to control diet ± standard error. See Additional File 6 for full results. E. Select differentially expressed (Benjamini-Hochberg adjusted p-value < 0.1) genes of interest in *B. thetaiotaomicron* within the cecum of vehicle-treated mice consuming the Western (WD) diet. Data are represented as log_2_ fold change relative to control diet ± standard error. See Additional File 6 for full results. For all analyses, n = 4.

In comparison to diet, ciprofloxacin treatment had a relatively minor impact on *A. muciniphila* gene expression (Additional File 5). This likely relates to the fact that on both diets, ciprofloxacin did not impact the relative abundance of *A. muciniphila* (Figure 3A). In total, ciprofloxacin significantly altered the expression of 14 and 26 genes on the control and Western diets, respectively (Additional File 5). On the control diet, *A. muciniphila* increased the expression of the molecular chaperone protein DnaK, which is known to play a role in stress responses (73-75). Additionally, we observed elevated expression of a MoxR family ATPase following ciprofloxacin treatment on this diet. Though these proteins are poorly characterized, other members of this family have been shown to regulate stress responses in other bacteria (76). On the Western diet, several genes related to tryptophan biosynthesis and metabolism were elevated following ciprofloxacin treatment (AMUC_RS08210 - tryptophan synthase subunit beta, AMUC_RS08190 - anthranilate synthase component I family protein, AMUC_RS08215 - tryptophan synthase subunit alpha); however, their biological significance is unclear at this time (Additional File 5). Lastly, an examination of the interaction between diet and ciprofloxacin treatment indicated that only six genes (two of which encoded tRNAs) were significantly altered, suggesting that diet does not have a major impact on the response of this bacterium to ciprofloxacin within the microbiome (Additional File 5).

In contrast to *A. muciniphila*, diet had a relatively minor impact on *B. thetaiotaomicron* gene expression while ciprofloxacin induced extensive changes. It is important to note that *B. thetaiotaomicron* bloomed in response to the Western diet and was significantly perturbed on the Western diet but not on the control (Figure 3C). In total, *B. thetaiotaomicron* altered the expression of 74 genes in response to Western diet consumption (Additional File 6). Of note, this diet increased the expression of an aminoglycoside efflux pump (BT_0305), the universal stress protein UspA (BT_0901), and a hemin receptor (BT_0316). However, more than half of the genes (52.7%) that changed in response to diet are of unknown function and are classified as “hypothetical proteins;” thus, it is difficult to draw conclusions without improved annotation. Overall, ciprofloxacin appeared to induce extensive transcriptional changes within *B. thetaiotaomicron* regardless of diet. On the control diet, we observed an increased abundance of transcripts encoding a number of proteins involved in capsular polysaccharide (CPS) biosynthesis and export (Figure 3D, Additional File 6). Within *B. thetaiotaomicron*, CPS production is encoded by a total of 182 genes distributed among eight loci (typically termed *cps1-8*) (77, 78). It is hypothesized that an individual bacterium expresses one of these CPS configurations at any given time and that these structures play key roles in processes such as nutrient acquisition and immune evasion (78). Additionally, the two genes with the greatest increase in expression during ciprofloxacin treatment encoded UDP-glucose 6-dehydrogenase, which plays a key role in the biosynthesis of glycan precursors that are essential for capsule production in other bacteria (79-81). Together, these findings may suggest a role for CPS state as a determinant of ciprofloxacin susceptibility *in vivo*.

On the Western diet, ciprofloxacin elicited profound changes in transcriptional activity, altering the expression of 442 different genes (Figure 3E, Additional File 6) and this robust response may be related to the reduction in *B. thetaiotaomicron* in this condition (Figure 3E). Interestingly, *B. thetaiotaomicron* reduced the expression of a number of genes involved in the utilization of host-derived carbohydrates (sialic acid-specific 9-O-acetylesterase, endo-beta-N-acetylglucosaminidase F1, beta-hexosaminidase) and stress responses (universal stress protein UspA, thioredoxin), mirroring the changes we saw at the whole-community level (Figure 3E, Additional File 6). Conversely, we observed increased expression of several genes that encode molecular chaperones (such as GroEL and GroES) or are involved in DNA replication or damage repair (such as a DNA helicase, DNA gyrase subunit B, DNA mismatch repair protein MutS, DNA polymerase III subunit alpha, Holliday junction DNA helicase RuvB, DNA-binding proteins, and DNA primase) (Figure 3E, Additional File 6). Ciprofloxacin, a fluoroquinolone class antimicrobial, triggers DNA damage via inhibition of DNA gyrase and topoisomerase IV. Thus, these changes in gene expression may be reflective of the primary mechanism of action of this antibiotic and are consistent with previously published data (20). Furthermore, our ability to detect such changes is an important indication that our analysis is detecting ciprofloxacin-induced transcriptional shifts. Lastly, diet appears to have a significant impact on ciprofloxacin-induced transcriptional changes within *B. thetaiotaomicron*, modulating the response of 148 genes (Additional File 6). Of note, Western diet consumption in the context of ciprofloxacin treatment had a negative impact on several genes involved in the acquisition of nutrients, such as vitamin B_12_ and hemin receptors, and transporters of glucose/galactose, hexuronate, arabinose, and Na^+^ (Additional File 6). Thus, it is likely that the availability of nutrients within the gut plays a role in the response of these bacteria to antibiotics.

## Discussion

Previous work has demonstrated that host diet, particularly with respect to sugar and fiber content, plays a major role in the extent of antibiotic-induced microbiome disruption (20). In Western societies, many people consume a diet high in added sugars and fat, but low in host-indigestible fiber. It is thought that such a composition promotes the development of metabolic syndrome, heart disease, diabetes, and a number of other chronic conditions (36-46). Furthermore, broad-spectrum antibiotic use and resulting microbiome dysbiosis have been associated with a number of similar co-morbidities along with increased susceptibility to opportunistic infections (1, 5-7, 21-23, 25, 26). Despite this connection, little work has been done examining how host dietary composition impacts the response of the microbiota to antibiotic perturbation. It is known that nutrient availability and metabolic state are a major determinant of antibiotic susceptibility of bacteria *in vitro* (20, 64-67, 82-88). Thus, it is likely that modulating host diet, thus changing the availability of nutrients to the microbiota, would alter the sensitivity of bacteria in these communities to antibiotic therapy.

To address this knowledge gap, we utilized a combined metagenomic and metatranscriptomic approach to profile taxonomic and functional changes in response to both diet and antibiotic treatment. By utilizing these tools in parallel, we are able to link transcriptional changes to observed shifts in community structure on each diet. Using metagenomics, we observed that ciprofloxacin had a differential impact on community composition that was diet-dependent. Specifically, we observed a statistically significant expansion of the *Firmicutes* phylum following ciprofloxacin treatment on the Western, but not control, diet. Using metatranscriptomics, we observed that ciprofloxacin treatment resulted in a decreased abundance of transcripts from the TCA cycle in both diets, suggesting that this response is diet-independent. Furthermore, this observation is consistent with previous *in vitro* findings that demonstrate a key role for bacterial respiration as a determinant of susceptibility to fluoroquinolones (64-66, 68, 82, 85). Conversely, ciprofloxacin had diverging impacts on the abundance of various iron and mucin utilization transcripts on the Western and control diets. Most notably, we found that Western diet consumption (alone and in the presence of ciprofloxacin) influenced the abundance of transcripts encoding genes known to play a role in virulence, supporting previous literature demonstrating that nutrient availability impacts virulence of enteric pathogens (20, 89-92). Lastly, we detected species-specific transcriptional changes in two important commensal bacteria, *B. thetaiotaomicron* and *A. muciniphila*. In addition to detecting changes in transcript levels that were reflective of stress responses, we also observed that transcripts involved in diverse cellular processes such as nutrient acquisition, carbon metabolism, and capsular polysaccharide (CPS) biosynthesis were differentially expressed as well.

Despite the advantages of a multi-omic approach, there are a number of drawbacks to these techniques that complicate the interpretation of our results. Most crucially, nearly all analytical pipelines used to analyze microbiome data are reliant on existing databases that are largely incomplete. It is hypothesized that approximately half of all genes within the human gut microbiome have no functional annotation (93). Thus, the ability to accurately profile the transcriptional activity of these communities is inherently limited by the quality and completeness of the databases utilized. Additionally, elucidating the biological significance of taxonomic of functional changes is often difficult in many microbiome analyses. Due to the complex nature of these communities, it is often difficult to ascertain if the observed transcriptional changes are the result of the direct action of the antibiotic, or the indirect effect of changes in host physiology, nutrient availability, or the disruption of ecological networks within the microbiome. For example, our transcriptional analysis of *B. thetaiotaomicron* showed that this bacterium differentially expressed receptors for both hemin and vitamin B_12_, which may suggest that these nutrients play a role in ciprofloxacin toxicity. Alternatively, it is possible that these transcriptional changes are reflective of increased availability of these nutrients due to decreased competition from other members of the microbiota (though these hypotheses are not mutually exclusive). Additionally, it is possible that dietary composition could play a significant role in antibiotic absorption or sequestration in the gut, which in turn would impact the extent of the damage caused to the microbiota.

This study builds on recent work that demonstrates that the availability of metabolites to the bacteria in the host plays an important role in determining the extent of antibiotic-induced microbiome disruption (20). Taken together, these results demonstrate the need to consider dietary composition in the design and interpretation of experiments focused on understanding the impact of antibiotics on the microbiota. Previous studies have demonstrated that dietary changes induce rapid shifts in gut microbiome composition (32, 34, 43, 56, 94-97). Therefore, in the long-term, dietary modulation could represent an attractive strategy to reduce the collateral damage to commensal bacteria and the resulting complications from dysbiosis caused by clinical therapy. Despite these promising applications, considerable work is required before these findings have direct clinical relevance. In particular, the considerable differences in physiology, microbiome composition, and diet between humans and rodents complicate the direct clinical relevance of these findings. Furthermore, it is unclear whether short-term dietary modulation has any long-term consequences on either the host or the microbiome. Thus, additional research is warranted to fully elucidate how host diet impacts antibiotic-induced microbiome disruption in humans.

## Conclusions

Using a combined metagenomic and metatranscriptomic approach, we demonstrate that murine diet composition has a major impact on the response of the murine gut microbiome to ciprofloxacin therapy. First, we found that the gut microbiome undergoes differential shifts in community structure in response to antibiotic treatment in a diet-dependent manner. At the transcriptional level, we found that ciprofloxacin reduced the abundance of TCA cycle transcripts regardless of diet, suggesting that central carbon metabolism plays a role in the activity of this antibiotic *in vivo*. Despite this commonality, we observed extensive differences in the transcriptional response of the microbiome to dietary intervention and/or ciprofloxacin treatment. Notably, mice consuming a Western diet had a high abundance of transcripts encoding proteins known to degrade host-derived polysaccharides such as sialic residues of mucin, suggesting that the consumption of this diet may have detrimental impacts on host physiology. Lastly, we identified species-specific changes in transcript abundance in two key members of the gut microbiome, *A. muciniphila* and *B. thetaiotaomicron*. In *A. muciniphila*, consumption of a Western diet increased the expression of several genes known to play a role in stress responses. In *B. thetaiotaomicron*, we found that a number of genes involved in CPS biosynthesis were differentially expressed during ciprofloxacin treatment on the control but not Western diet, which may suggest a divergent response of this bacterium to ciprofloxacin that is dependent on nutrient composition. Taken together, these findings demonstrate that host diet is a determinant of antibiotic-induced microbiome perturbations.

## Methods

### Animal Procedures

All animal work was approved by Brown University’s Institutional Animal Care and Use Committee (IACUC) under protocol number 1706000283. 4-week-old female C57BL/6 mice were purchased from Jackson Laboratories (Bar Harbor, ME, USA) and given a 2-week habituation period immediately following arrival at Brown University’s Animal Care Facility. After habituation, mice were switched from standard chow (Laboratory Rodent Diet 5001, St. Louis, MO, USA) to either a Western diet (D12079B, Research Diets Inc., New Brunswick, NJ, USA) or a macronutrient-defined control diet (D122405B, Research Diets Inc., New Brunswick, NJ, USA) for 1 week. Following dietary intervention, mice were given acidified ciprofloxacin (12.5 mg/kg/day), or a pH-adjusted vehicle, via filter-sterilized drinking water *ad libitum* for 24 hours (n = 8-12 per treatment group). Water consumption was monitored to assure equal consumption across cages. Mice were then sacrificed and dissected in order to collect cecal contents. Cecal contents were immediately transferred to ZymoBIOMICS DNA/RNA Miniprep Kit (Zymo Research, Irvine, CA, USA) Collection Tubes containing DNA/RNA Shield. Tubes were processed via vortex at maximum speed for 5 minutes to homogenize cecal contents, then placed on ice until permanent storage at −80°C.

### Nucleic Acid Extraction & Purification

Total nucleic acids (DNA and RNA) were extracted from samples using the ZymoBIOMICS DNA/RNA Miniprep Kit from Zymo Research (R2002, Irvine, CA, USA) using the parallel extraction protocol as per the manufacturer instructions. Total RNA and DNA were eluted in nuclease-free water and quantified using the dsDNA-HS and RNA-HS kits on a Qubit™ 3.0 fluorometer (Thermo Fisher Scientific, Waltham, MA, USA) before use in library preparations.

### 16S rRNA Amplicon Preparation & Sequencing

The 16S rRNA V4 hypervariable region was amplified from total DNA using the barcoded 518F forward primer and the 816Rb reverse primers from the Earth Microbiome Project (98). Amplicons were generated using 5X Phusion High-Fidelity DNA Polymerase under the following cycling conditions: initial denaturation at 98°C for 30 seconds, followed by 25 cycles of 98°C for 10 seconds, 57°C for 30 seconds, and 72°C for 30 seconds, then a final extension at 72°C for 5 minutes. After amplification, samples were pooled in equimolar amounts and visualized via gel electrophoresis. Pooled amplicon library was submitted to the Rhode Island Genomics and Sequencing Center at the University of Rhode Island (Kingston, RI, USA) for sequencing on the Illumina MiSeq platform. Amplicons were pair-end sequenced (2×250 bp) using the 500-cycle kit with standard protocols. We obtained an average of 106,135 ± 49,789 reads per sample. Raw reads were deposited in the NCBI Short Read Archive (SRA) under accession number PRJNA594642.

### Analysis of 16S rRNA Sequencing Reads

Raw 16S rRNA reads were subjected to quality filtering, trimming, de-noising, and merging using the DADA2 package (version 1.8.0) in R (version 3.5.0). Ribosomal sequence variants were assigned taxonomy using the RDP Classifier algorithm with RDP Training set 16 using the *assignTaxonomy* function in DADA2 (99). Alpha diversity (Shannon) and beta diversity (Bray-Curtis dissimilarity) were calculated using the phyloseq package (version 1.24.2) in R (version 3.5.0).

### Metagenomic & Metatranscriptomic Library Preparation

Metagenomic libraries were prepared from DNA (100 ng) using the NEBNext® Ultra II FS DNA Library Prep Kit (New England BioLabs, Ipswich, MA, USA) > 100ng input protocol as per the manufacturer’s instructions. This yielded a pool of 200 – 1000 bp fragments where the average library was 250-500 bp. Metatranscriptomic libraries were prepared from total RNA using the NEBNext® Ultra II Directional RNA Sequencing Prep Kit (New England BioLabs, Ipswich, MA, USA) in conjunction with the NEBNext® rRNA Depletion Kit for Human/Mouse/Rat (New England BioLabs, Ipswich, MA, USA) and the MICROBExpress kit (Invitrogen, Carlsbad, CA, USA). First, up to 1 ug of total RNA was treated with rDNase I and subsequently depleted of bacterial rRNAs using MICROBExpress as per the manufacturer’s instructions. This depleted RNA was then used to prepare libraries with the NEBNext® Ultra II Directional RNA Sequencing Prep & rRNA depletion kits as per the manufacturer’s instructions. This yielded libraries that averaged between 200-450 bp. Once library preparation was complete, both metagenomic and metatranscriptomic libraries were sequenced as paired-end 150 bp reads on an Illumina HiSeq X Ten. We sequenced an average of 2,278,948,631 (± 2,309,494,556) bases per metagenomic sample and 14,751,606,319 (± 3,089,205,166) bases per metatranscriptomic sample. One metagenomic sample from the Western diet + vehicle group had an abnormally low number of bases sequenced (165,000 bp) and was excluded from all subsequent analyses. Following the removal of this sample, we obtained an average of 2,430,867,540 (± 2,306,317,898) bases per metagenomic sample. All reads were deposited in the NCBI Short Read Archive under BioProject number PRJNA563913.

### Processing of Raw Metagenomic and Metatranscriptomic Reads

Raw metagenomic reads were trimmed and decontaminated using kneaddata utility (version 0.6.1) (100). In brief, reads were first trimmed to remove low quality bases and Illumina TruSeq3 adapter sequences using trimmomatic (version 0.36) using SLIDINGWINDOW value of 4:20 and ILLUMINACLIP value 2:20:10, respectively (101). Trimmed reads shorter than 75 bases were discarded. Reads passing quality control were subsequently decontaminated by removing those that mapped to the genome of C57BL/6J mice using bowtie2 (version 2.2) (102). Additionally, preliminary work by our group detected high levels of reads mapping to two murine retroviruses found in our animal facility: murine mammary tumor virus (MMTV) and murine osteosarcoma viruses (MOV) (20). Raw metatranscriptomic reads were trimmed and decontaminated using the same parameters. However, in addition to removing reads that mapped to the C57BL/6J, MMTV, and MOV genomes, we also decontaminated sequences that aligned to the SILVA 128 LSU and SSU Parc ribosomal RNA databases (103).

### Taxonomic Classification of Metagenomic Reads

Trimmed and decontaminated metagenomic reads were taxonomically classified against a database containing all bacterial and archaeal genomes found in NCBI RefSeq using Kraken2 (version 2.0.7-beta) with a default k-mer length of 35 (104). Phylum- and species-level abundances were subsequently calculated from Kraken2 reports using Bracken (version 2.0.0) with default settings (105). The phyloseq package (version 1.28.0) in R (version 3.6.0) was used to calculate alpha diversity using the Shannon Diversity Index (106). Metagenomic data was not subsampled prior to analysis.

To perform differential abundance testing, species-level taxonomic output was first filtered to remove taxa that were not observed in >1,000 reads (corresponding to approximately 0.1% of all reads) in at least 20% of all samples using phyloseq in R. Differential abundance testing was subsequently performed on filtered counts using the DESeq2 package (version 1.24.0) using default parameters (62). All p-values were corrected for multiple hypothesis testing using the Benjamini-Hochberg method (107).

### Annotation of Metatranscriptomic Reads Using SAMSA2

Trimmed and decontaminated metatranscriptomic reads were annotated using a modified version of the Simple Annotation of Metatranscriptomes by Sequence Analysis 2 (SAMSA2) pipeline as described previously (20, 69, 108). First, the Paired-End Read Merger (PEAR) utility was used to merge forward and reverse reads (109). Merged reads were then aligned to databases containing entries from the RefSeq and SEED Subsystems databases using DIAMOND (version 0.9.12) (110, 111). The resulting alignment counts were subsequently analyzed using DESeq2 (version 1.24.0) using the Benjamini-Hochberg method to perform multiple hypothesis testing correction (20, 69, 107). Features with an adjusted p-value of less than 0.1 were considered to be statistically significant.

### Metatranscriptomic Analysis using HUMAnN2

To determine the impact of dietary modulation and ciprofloxacin treatment on gene expression within the gut microbiome, we used the HMP Unified Metabolic Analysis Network 2 (HUMAnN2, version 0.11.1) pipeline (63). First, metagenomic reads were taxonomically annotated using MetaPhlan2 (version 2.6.0) and functionally annotated against the UniRef90 database to generate gene family and MetaCyc pathway level abundances. To ensure consistent assignment between paired samples, the taxonomic profile generated from the metagenomic reads was supplied to the HUMAnN2 algorithm during the analysis of the corresponding metatranscriptomic reads. Metatranscriptomic reads were subsequently annotated as done for metagenomic reads. The resulting gene family and pathway level abundance data from the metatranscriptomic reads was normalized against the metagenomic data from the corresponding sample and smoothed using the Witten-Bell method (112). Lastly, the resulting RPKM values were unstratified to obtain whole-community level data, converted into relative abundances, and analyzed using LEfSe (version 1) hosted on the Galaxy web server (113).

### Transcriptional Analysis of *A. muciniphila* and *B. thetaiotaomicron*

A modified version of a previously published pipeline from Deng *et al.* was utilized to perform transcriptional analysis of individual species within the murine microbiome during dietary modulation and antibiotic treatment (20, 71). First, Kraken2 (version 2.0.7-beta) was used to identify the fifty most prevalent bacterial species present within the metatranscriptomic samples (104). Next, the BBSplit utility within the BBMap package (version 37.96) was used to extract reads within our metatranscriptomic dataset that mapped to these fifty most abundant species (114). Reads from *B. thetaiotaomicron* and *A. muciniphila* were subsequently aligned to their corresponding reference genomes using the BWA-MEM algorithm (version 0.7.15) (115). Lastly, the featureCounts command within the subread program (version 1.6.2) was used to analyze the resulting alignment files to generate a count table for differential expression analysis with DESeq2 (62). All p-values were corrected for multiple hypothesis testing with the Benjamini-Hochberg method (107). Features with an adjusted p-value of less than 0.1 were considered to be statistically significant.

## List of Abbreviations

PTS: phosphotransferase systems
MACs: microbiota-accessible carbohydrates
SCFAs: short chain fatty acids
TCA: tricarboxylic acid
HUMAnN2: Human Microbiome Project (HMP) Unified Metabolic Analysis Network 2
CPS: capsular polysaccharide
GH98: glycoside hydrolase family 98
SAMSA2: Simple Annotation of Metatranscriptomes by Sequence Analysis 2
SRA: Sequence Read Archive

## Declarations

### Availability of Data and Materials

The datasets generated and analyzed during this study are available from the NCBI Short Read Archive (SRA) under BioProject accession numbers PRJNA563913 (metagenomics and metatranscriptomics) and PRJNA594642 (16S rRNA amplicon sequences). Any additional information is available from the corresponding author upon request.

### Authors’ contributions

DJC planned the study, performed mouse experiments, extracted nucleic acids from cecal samples, conducted analysis of 16S rRNA amplicon, metagenomic, and metatranscriptomic data, and co-wrote the manuscript. JIW assisted with mouse experiments, prepared DNA and RNA into sequencing libraries for metagenomics and metatranscriptomics, assisted in the interpretation of results, and co-wrote the manuscript. BJK assisted with the analysis of metatranscriptomic data. SP assisted in the interpretation of results. PB conceptualized and planned the study, contributed to the writing of the manuscript, and secured funding. All authors have read and approved of the final manuscript.

## Acknowledgements

D.J.C, B.J.K, and J.I.W., and S.P. were supported by the Graduate Research Fellowship Program from the National Science Foundation under award number 1644760. P.B. was supported by the National Center for Complementary & Integrative Health of the NIH Award Number R21AT010366, and institutional development awards P20GM121344 and P20GM109035 received from the National Institute of General Medical Sciences within the NIH. The funding agencies had no role in the design of the study or the collection, analysis, and interpretation of data.

## Figures

**Figure S1:**
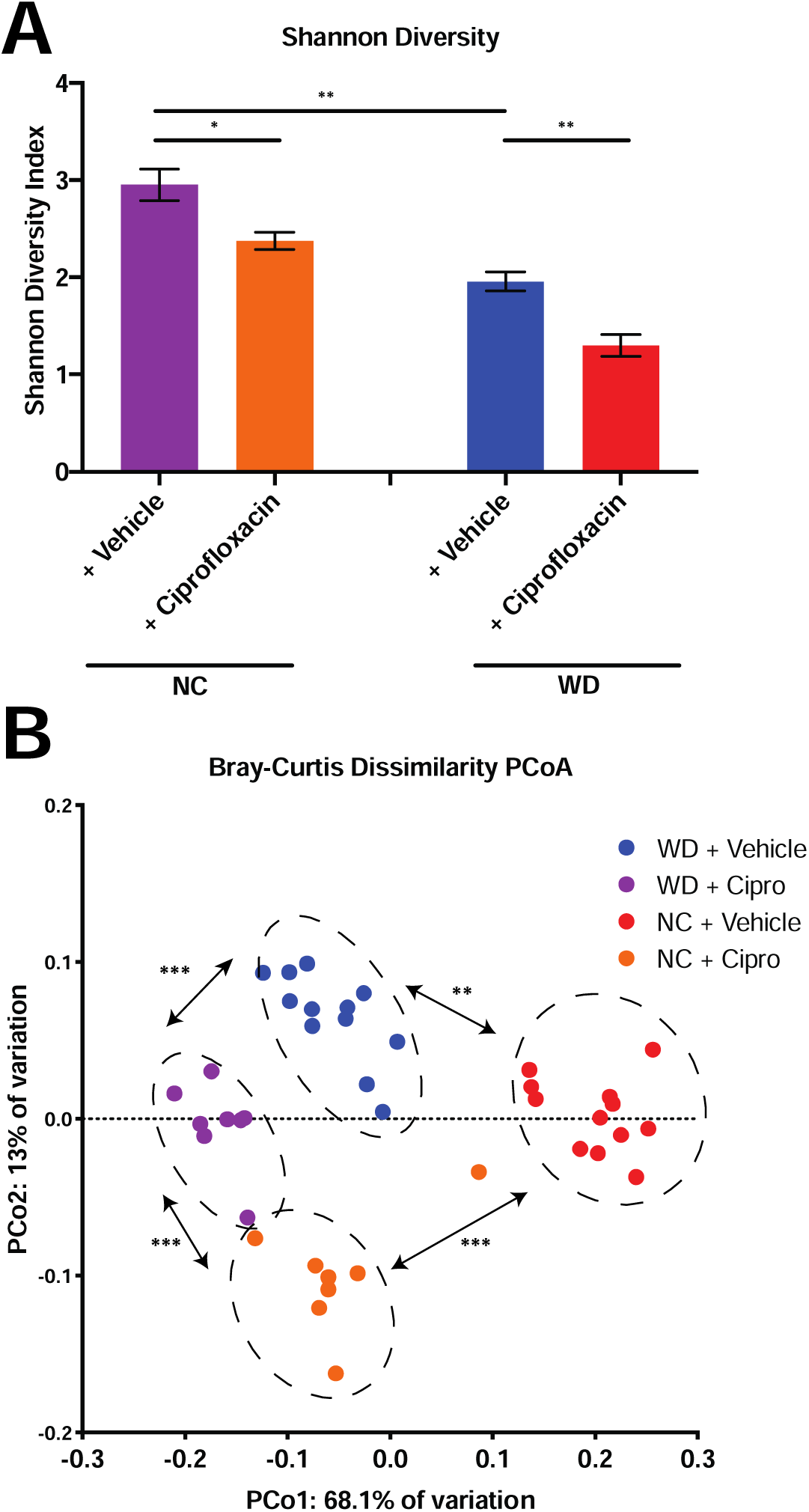
Dietary composition and antibiotic treatment impact the diversity of the gut microbiome. A. Alpha diversity of experimental groups as measured by the Shannon Diversity Index. Data are represented as mean ± standard error of the mean (SEM). (*p<0.05, **p<0.01, ***p<0.001, Welch ANOVA with Dunnett T3 test for multiple hypothesis testing) B. Principle coordinate analysis of experimental groups as measured by Bray-Curtis dissimilarity. (**p<0.01, ***p<0.001, Permutational ANOVA).

## Additional Files

**Additional File 1:** Full DESeq2 results of differential abundance testing of top 90 species detected by shotgun metagenomics

Table S1 – Differential abundance testing of the impact of Western diet (WD) consumption on the abundance of the top 90 bacterial species detected in our dataset. Log_2_ fold change values were calculated relative to control diet samples.

Table S2 – Differential abundance testing of the impact of ciprofloxacin treatment on the abundance of the top 90 bacterial species in mice consuming the Western diet (WD). Log_2_ fold change values were calculated relative to vehicle-treated samples on the WD.

Table S3 - Differential abundance testing of the impact of ciprofloxacin treatment on the abundance of the top 90 bacterial species in mice consuming the control diet (NC). Log_2_ fold change values were calculated relative to vehicle-treated samples on the NC.

Table S4 – Interaction term analysis generated by DESeq2 for the impact of host diet consumption on changes in species abundance following ciprofloxacin therapy. Log_2_ fold change values were calculated relative to vehicle-treated samples on the NC.

**Additional File 2:** Full LEfSe results from the analysis of MetaCyc pathway abundance generated by HUMAnN2. “Class” denotes the experimental group a particular pathway was associated with.

Table S5 – LEfSe analysis of all experimental groups.

Table S6 – Pairwise LEfSe analysis of vehicle-treated samples from mice consuming either the Western (WD) or control (NC) diet.

Table S7– Pairwise LEfSe analysis of ciprofloxacin- and vehicle-treated samples from mice consuming the control diet (NC)

Table S8 – Pairwise LEfSe analysis of ciprofloxacin- and vehicle-treated samples from mice consuming the Western diet (WD)

**Additional File 3:** Full DESeq2 results of SEED subsystem abundance generated by SAMSA2

Table S9 – Differential abundance testing of the impact of Western diet (WD) consumption on the abundance of SEED subsystems in the murine cecal metatranscriptome. Log_2_ fold change values were calculated relative to control diet samples.

Table S10 – Differential abundance testing of the impact of ciprofloxacin treatment on the abundance of SEED subsystems in the murine cecal metatranscriptome in animals consuming the Western diet (WD). Log_2_ fold change values were calculated relative to vehicle-treated samples on the WD.

Table S11 - Differential abundance testing of the impact of ciprofloxacin treatment on the abundance of SEED subsystems in the murine cecal metatranscriptome in animals consuming the control diet (NC). Log_2_ fold change values were calculated relative to vehicle-treated samples on the NC.

**Additional File 4:** Full DESeq2 results of RefSeq transcript abundance generated by SAMSA2

Table S12 – Differential abundance testing of the impact of Western diet (WD) consumption on the abundance of RefSeq transcripts in the murine cecal metatranscriptome. Log_2_ fold change values were calculated relative to control diet samples.

Table S13 – Differential abundance testing of the impact of ciprofloxacin treatment on the abundance of RefSeq transcripts in the murine cecal metatranscriptome in animals consuming the Western diet (WD). Log_2_ fold change values were calculated relative to vehicle-treated samples on the WD.

Table S14 - Differential abundance testing of the impact of ciprofloxacin treatment on the abundance of RefSeq transcripts in the murine cecal metatranscriptome in animals consuming the control diet (NC). Log_2_ fold change values were calculated relative to vehicle-treated samples on the NC.

Table S15 – Interaction term analysis generated by DESeq2 for the impact of host diet consumption on changes in RefSeq transcripts abundance following ciprofloxacin therapy. Log_2_ fold change values were calculated relative to vehicle-treated samples on the NC.

**Additional File 5:** Full DESeq2 results of transcript abundance analysis of *A. muciniphila* during dietary intervention and ciprofloxacin treatment

Table S16 – Differential abundance testing of the impact of Western diet (WD) consumption on the abundance of *A. muciniphila* transcripts within the murine cecal metatranscriptome. Log_2_ fold change values were calculated relative to control diet samples.

Table S17 – Differential abundance testing of the impact of ciprofloxacin treatment on the abundance of *A. muciniphila* transcripts within the murine cecal metatranscriptome in animals consuming the Western diet (WD). Log_2_ fold change values were calculated relative to vehicle-treated samples on the WD.

Table S18 - Differential abundance testing of the impact of ciprofloxacin treatment on the abundance of *A. muciniphila* transcripts within the murine cecal metatranscriptome in animals consuming the control diet (NC). Log_2_ fold change values were calculated relative to vehicle-treated samples on the NC.

Table S19 – Interaction term analysis generated by DESeq2 for the impact of host diet consumption on changes in *A. muciniphila* transcript abundance following ciprofloxacin therapy. Log_2_ fold change values were calculated relative to vehicle-treated samples on the NC.

**Additional File 6:** Full DESeq2 results of transcript abundance analysis of *B. thetaiotaomicron* during dietary intervention and ciprofloxacin treatment

Table S20 – Differential abundance testing of the impact of Western diet (WD) consumption on the abundance of *B. thetaiotaomicron* transcripts within the murine cecal metatranscriptome. Log_2_ fold change values were calculated relative to control diet samples.

Table S21 – Differential abundance testing of the impact of ciprofloxacin treatment on the abundance of *B. thetaiotaomicron* transcripts within the murine cecal metatranscriptome in animals consuming the Western diet (WD). Log_2_ fold change values were calculated relative to vehicle-treated samples on the WD.

Table S22 - Differential abundance testing of the impact of ciprofloxacin treatment on the abundance of *B. thetaiotaomicron* transcripts within the murine cecal metatranscriptome in animals consuming the control diet (NC). Log_2_ fold change values were calculated relative to vehicle-treated samples on the NC.

Table S23 – Interaction term analysis generated by DESeq2 for the impact of host diet consumption on changes in *B. thetaiotaomicron* transcript abundance following ciprofloxacin therapy. Log_2_ fold change values were calculated relative to vehicle-treated samples on the NC.

## References

1. Rowan-Nash AD, Korry BJ, Mylonakis E, Belenky P. 2019. Cross-Domain and Viral Interactions in the Microbiome. Microbiol Mol Biol Rev 83:51.

2. Gilbert JA, Blaser MJ, Caporaso JG, Jansson JK, Lynch SV, Knight R. 2018. Current understanding of the human microbiome. Nat Med 24:392–400.

3. Ursell LK, Metcalf JL, Parfrey LW, Knight R. 2012. Defining the human microbiome. Nutr Rev 70:S38–44.

4. Stiemsma LT, Michels KB. 2018. The Role of the Microbiome in the Developmental Origins of Health and Disease. Pediatrics 141:e20172437.

5. Blaser M. 2011. Antibiotic overuse: Stop the killing of beneficial bacteria. Nature 476:393–394.

6. Mukherjee PK, Chandra J, Retuerto M, Sikaroodi M, Brown RE, Jurevic R, Salata RA, Lederman MM, Gillevet PM, Ghannoum MA. 2014. Oral mycobiome analysis of HIV-infected patients: identification of Pichia as an antagonist of opportunistic fungi. PLoS Pathog 10:e1003996.

7. Peleg AY, Hogan DA, Mylonakis E. 2010. Medically important bacterial-fungal interactions. Nat Rev Microbiol 8:340–349.

8. De Luca F, Shoenfeld Y. 2018. The microbiome in autoimmune diseases. Clin Exp Immunol 195:74–85.

9. Dickerson F, Severance E, Yolken R. 2017. The microbiome, immunity, and schizophrenia and bipolar disorder. Brain Behav Immun 62:46–52.

10. Foster JA, McVey Neufeld K-A. 2013. Gut-brain axis: how the microbiome influences anxiety and depression. Trends Neurosci 36:305–312.

11. Foster JA, Rinaman L, Cryan JF. 2017. Stress & the gut-brain axis: Regulation by the microbiome. Neurobiology of Stress 7:124–136.

12. Hartstra AV, Bouter KEC, Bäckhed F, Nieuwdorp M. 2015. Insights into the role of the microbiome in obesity and type 2 diabetes. Diabetes Care 38:159–165.

13. Leong KSW, Derraik JGB, Hofman PL, Cutfield WS. 2018. Antibiotics, gut microbiome and obesity. Clin Endocrinol (Oxf) 88:185–200.

14. Lynch SV, Boushey HA. 2016. The microbiome and development of allergic disease. Curr Opin Allergy Clin Immunol 16:165–171.

15. Riiser A. 2015. The human microbiome, asthma, and allergy. Allergy Asthma Clin Immunol 11:35.

16. Rogers MB, Firek B, Shi M, Yeh A, Brower-Sinning R, Aveson V, Kohl BL, Fabio A, Carcillo JA, Morowitz MJ. 2016. Disruption of the microbiota across multiple body sites in critically ill children. Microbiome 4:66.

17. Tremlett H, Bauer KC, Appel-Cresswell S, Finlay BB, Waubant E. 2017. The gut microbiome in human neurological disease: A review. Ann Neurol 81:369–382.

18. Vieira SM, Pagovich OE, Kriegel MA. 2014. Diet, microbiota and autoimmune diseases. Lupus 23:518–526.

19. Dethlefsen L, Relman DA. 2011. Incomplete recovery and individualized responses of the human distal gut microbiota to repeated antibiotic perturbation. Proc Natl Acad Sci USA 108 Suppl 1:4554–4561.

20. Cabral DJ, Penumutchu S, Reinhart EM, Zhang C, Korry BJ, Wurster JI, Nilson R, Guang A, Sano WH, Rowan-Nash AD, Li H, Belenky P. 2019. Microbial Metabolism Modulates Antibiotic Susceptibility within the Murine Gut Microbiome. Cell Metab 30:800–823.

21. Chang JY, Antonopoulos DA, Kalra A, Tonelli A, Khalife WT, Schmidt TM, Young VB. 2008. Decreased diversity of the fecal Microbiome in recurrent Clostridium difficile-associated diarrhea. J Infect Dis 197:435–438.

22. Preidis GA, Versalovic J. 2009. Targeting the human microbiome with antibiotics, probiotics, and prebiotics: gastroenterology enters the metagenomics era. Gastroenterology 136:2015–2031.

23. Theriot CM, Bowman AA, Young VB. 2016. Antibiotic-Induced Alterations of the Gut Microbiota Alter Secondary Bile Acid Production and Allow for Clostridium difficile Spore Germination and Outgrowth in the Large Intestine. mSphere 1:e00045–15.

24. Rea MC, Dobson A, O’Sullivan O, Crispie F, Fouhy F, Cotter PD, Shanahan F, Kiely B, Hill C, Ross RP. 2011. Effect of broad- and narrow-spectrum antimicrobials on Clostridium difficile and microbial diversity in a model of the distal colon. Proc Natl Acad Sci USA 108:4639–4644.

25. Rafii F, Sutherland JB, Cerniglia CE. 2008. Effects of treatment with antimicrobial agents on the human colonic microflora. Ther Clin Risk Manag 4:1343–1358.

26. Lessa FC, Winston LG, McDonald LC, Emerging Infections Program C. difficile Surveillance Team. 2015. Burden of Clostridium difficile infection in the United States. N Engl J Med 372:2369–2370.

27. Hryckowian AJ, Van Treuren W, Smits SA, Davis NM, Gardner JO, Bouley DM, Sonnenburg JL. 2018. Microbiota-accessible carbohydrates suppress Clostridium difficile infection in a murine model. Nature Microbiology 3:662–669.

28. Durack J, Huang YJ, Nariya S, Christian LS, Mark Ansel K, Beigelman A, Castro M, Dyer A-M, Israel E, Kraft M, Martin RJ, Mauger DT, Rosenberg SR, King TS, White SR, Denlinger LC, Holguin F, Lazarus SC, Lugogo N, Peters SP, Smith LJ, Wechsler ME, Lynch SV, Boushey HA, National Heart, Lung and Blood Institute’s “AsthmaNet.” 2018. Bacterial biogeography of adult airways in atopic asthma. Microbiome 6:104.

29. Wu GD, Chen J, Hoffmann C, Bittinger K, Chen Y-Y, Keilbaugh SA, Bewtra M, Knights D, Walters WA, Knight R, Sinha R, Gilroy E, Gupta K, Baldassano R, Nessel L, Li H, Bushman FD, Lewis JD. 2011. Linking long-term dietary patterns with gut microbial enterotypes. Science 334:105–108.

30. Smits SA, Leach J, Sonnenburg ED, Gonzalez CG, Lichtman JS, Reid G, Knight R, Manjurano A, Changalucha J, Elias JE, Dominguez-Bello MG, Sonnenburg JL. 2017. Seasonal cycling in the gut microbiome of the Hadza hunter-gatherers of Tanzania. Science 357:802–806.

31. Smits SA, Marcobal A, Higginbottom S, Sonnenburg JL, Kashyap PC. 2016. Individualized Responses of Gut Microbiota to Dietary Intervention Modeled in Humanized Mice. mSystems 1:105.

32. Bisanz JE, Upadhyay V, Turnbaugh JA, Ly K, Turnbaugh PJ. 2019. Meta-Analysis Reveals Reproducible Gut Microbiome Alterations in Response to a High-Fat Diet. Cell Host Microbe 26:265–272.

33. Ley RE, Bäckhed F, Turnbaugh P, Lozupone CA, Knight RD, Gordon JI. 2005. Obesity alters gut microbial ecology. Proc Natl Acad Sci USA 102:11070–11075.

34. Turnbaugh PJ, Ridaura VK, Faith JJ, Rey FE, Knight R, Gordon JI. 2009. The effect of diet on the human gut microbiome: a metagenomic analysis in humanized gnotobiotic mice. Science Translational Medicine 1:6ra14–6ra14.

35. Xu Z, Knight R. 2015. Dietary effects on human gut microbiome diversity. Br J Nutr 113 Suppl:S1–5.

36. Argueta DA, DiPatrizio NV. 2017. Peripheral endocannabinoid signaling controls hyperphagia in western diet-induced obesity. Physiology & Behavior 171:32–39.

37. Kanoski SE, Hsu TM, Pennell S. 2014. Obesity, Western Diet Intake, and Cognitive Impairment, pp. 57–62. In Omega-3 Fatty Acids in Brain and Neurological Health. Elsevier.

38. Sami W, Ansari T, Butt NS, Hamid MRA. 2017. Effect of diet on type 2 diabetes mellitus: A review. Int J Health Sci (Qassim) 11:65–71.

39. Qi L, Cornelis MC, Zhang C, van Dam RM, Hu FB. 2009. Genetic predisposition, Western dietary pattern, and the risk of type 2 diabetes in men. Am J Clin Nutr 89:1453–1458.

40. Sonnenburg ED, Sonnenburg JL. 2014. Starving our microbial self: the deleterious consequences of a diet deficient in microbiota-accessible carbohydrates. Cell Metab 20:779–786.

41. Sonnenburg ED, Sonnenburg JL. 2019. The ancestral and industrialized gut microbiota and implications for human health. Nat Rev Microbiol 17:383–390.

42. Sonnenburg JL, Xu J, Leip DD, Chen CH, Westover BP, Weatherford J, Buhler JD, Gordon JI. 2005. Glycan foraging in vivo by an intestine-adapted bacterial symbiont. Science 307:1955–1959.

43. Turnbaugh PJ. 2017. Microbes and Diet-Induced Obesity: Fast, Cheap, and Out of Control. Cell Host Microbe 21:278–281.

44. Trompette A, Gollwitzer ES, Yadava K, Sichelstiel AK, Sprenger N, Ngom-Bru C, Blanchard C, Junt T, Nicod LP, Harris NL, Marsland BJ. 2014. Gut microbiota metabolism of dietary fiber influences allergic airway disease and hematopoiesis. Nat Med 20:159–166.

45. Arpaia N, Campbell C, Fan X, Dikiy S, van der Veeken J, deRoos P, Liu H, Cross JR, Pfeffer K, Coffer PJ, Rudensky AY. 2013. Metabolites produced by commensal bacteria promote peripheral regulatory T-cell generation. Nature 504:451–455.

46. Cotillard A, Kennedy SP, Kong LC, Prifti E, Pons N, Le Chatelier E, Almeida M, Quinquis B, Levenez F, Galleron N, Gougis S, Rizkalla S, Batto J-M, Renault P, Doré J, Zucker J-D, Clement K, Ehrlich SD, Consortium AM. 2013. Dietary intervention impact on gut microbial gene richness. Nature 500:585–588.

47. Walker AW, Ince J, Duncan SH, Webster LM, Holtrop G, Ze X, Brown D, Stares MD, Scott P, Bergerat A, Louis P, McIntosh F, Johnstone AM, Lobley GE, Parkhill J, Flint HJ. 2011. Dominant and diet-responsive groups of bacteria within the human colonic microbiota. ISME J 5:220–230.

48. Fischbach MA, Sonnenburg JL. 2011. Eating for two: how metabolism establishes interspecies interactions in the gut. Cell Host Microbe 10:336–347.

49. Kashyap PC, Marcobal A, Ursell LK, Smits SA, Sonnenburg ED, Costello EK, Higginbottom SK, Domino SE, Holmes SP, Relman DA, Knight R, Gordon JI, Sonnenburg JL. 2013. Genetically dictated change in host mucus carbohydrate landscape exerts a diet-dependent effect on the gut microbiota. Proc Natl Acad Sci USA 110:17059–17064.

50. Yatsunenko T, Rey FE, Manary MJ, Trehan I, Dominguez-Bello MG, Contreras M, Magris M, Hidalgo G, Baldassano RN, Anokhin AP, Heath AC, Warner B, Reeder J, Kuczynski J, Caporaso JG, Lozupone CA, Lauber C, Clemente JC, Knights D, Knight R, Gordon JI. 2012. Human gut microbiome viewed across age and geography. Nature 486:222–227.

51. Wong J, de Souza R, Kendall C, Emam A, Jenkins D. 2006. Colonic health: Fermentation and short chain fatty acids. Journal of Clinical Gastroenterology 40:235–243.

52. Topping DL, Clifton PM. 2001. Short-chain fatty acids and human colonic function: Roles of resistant starch and nonstarch polysaccharides. Physiol Rev 81:1031–1064.

53. Macfarlane S, Macfarlane GT. 2003. Regulation of short-chain fatty acid production. Proceedings of the Nutrition Society 62:67–72.

54. Cani PD, Van Hul M, Lefort C, Depommier C, Rastelli M, Everard A. 2019. Microbial regulation of organismal energy homeostasis. Nat Metab 1:34–46.

55. Chambers ES, Preston T, Frost G, Morrison DJ. 2018. Role of Gut Microbiota-Generated Short-Chain Fatty Acids in Metabolic and Cardiovascular Health. Curr Nutr Rep 7:198–206.

56. David LA, Maurice CF, Carmody RN, Gootenberg DB, Button JE, Wolfe BE, Ling AV, Devlin AS, Varma Y, Fischbach MA, Biddinger SB, Dutton RJ, Turnbaugh PJ. 2014. Diet rapidly and reproducibly alters the human gut microbiome. Nature 505:559–563.

57. Poretsky R, Rodriguez-R LM, Luo C, Tsementzi D, Konstantinidis KT. 2014. Strengths and limitations of 16S rRNA gene amplicon sequencing in revealing temporal microbial community dynamics. PLoS ONE 9:e93827.

58. Ranjan R, Rani A, Metwally A, McGee HS, Perkins DL. 2016. Analysis of the microbiome: Advantages of whole genome shotgun versus 16S amplicon sequencing. Biochem Biophys Res Commun 469:967–977.

59. Clooney AG, Fouhy F, Sleator RD, Driscoll AO, Stanton C, Cotter PD, Claesson MJ. 2016. Comparing Apples and Oranges?: Next Generation Sequencing and Its Impact on Microbiome Analysis. PLoS ONE 11.

60. Tessler M, Neumann JS, Afshinnekoo E, Pineda M, Hersch R, Velho LFM, Segovia BT, Lansac-Toha FA, Lemke M, DeSalle R, Mason CE, Brugler MR. 2017. Large-scale differences in microbial biodiversity discovery between 16S amplicon and shotgun sequencing. Sci Rep 7:–14.

61. Shin N-R, Whon TW, Bae J-W. 2015. Proteobacteria: microbial signature of dysbiosis in gut microbiota. Trends Biotechnol 33:496–503.

62. Love MI, Huber W, Anders S. 2014. Moderated estimation of fold change and dispersion for RNA-seq data with DESeq2. Genome Biol 15:550.

63. Franzosa EA, McIver LJ, Rahnavard G, Thompson LR, Schirmer M, Weingart G, Lipson KS, Knight R, Caporaso JG, Segata N, Huttenhower C. 2018. Species-level functional profiling of metagenomes and metatranscriptomes. Nat Methods 15:962–968.

64. Meylan S, Porter CBM, Yang JH, Belenky P, Gutierrez A, Lobritz MA, Park J, Kim SH, Moskowitz SM, Collins JJ. 2017. Carbon Sources Tune Antibiotic Susceptibility in Pseudomonas aeruginosa via Tricarboxylic Acid Cycle Control. Cell Chem Biol 24:195–206.

65. Belenky P, Ye JD, Porter CBM, Cohen NR, Lobritz MA, Ferrante T, Jain S, Korry BJ, Schwarz EG, Walker GC, Collins JJ. 2015. Bactericidal Antibiotics Induce Toxic Metabolic Perturbations that Lead to Cellular Damage. Cell Rep 13:968–980.

66. Lobritz MA, Belenky P, Porter CBM, Gutierrez A, Yang JH, Schwarz EG, Dwyer DJ, Khalil AS, Collins JJ. 2015. Antibiotic efficacy is linked to bacterial cellular respiration. Proc Natl Acad Sci USA 112:8173–8180.

67. Dwyer DJ, Belenky PA, Yang JH, MacDonald IC, Martell JD, Takahashi N, Chan CTY, Lobritz MA, Braff D, Schwarz EG, Ye JD, Pati M, Vercruysse M, Ralifo PS, Allison KR, Khalil AS, Ting AY, Walker GC, Collins JJ. 2014. Antibiotics induce redox-related physiological alterations as part of their lethality. Proc Natl Acad Sci USA 111:E2100–9.

68. Dwyer DJ, Kohanski MA, Hayete B, Collins JJ. 2007. Gyrase inhibitors induce an oxidative damage cellular death pathway in Escherichia coli. Mol Syst Biol 3:91.

69. Westreich ST, Treiber ML, Mills DA, Korf I, Lemay DG. 2018. SAMSA2: a standalone metatranscriptome analysis pipeline. BMC Bioinformatics 19:175.

70. Corfield AP, Wagner SA, Clamp JR, Kriaris MS, Hoskins LC. 1992. Mucin degradation in the human colon: production of sialidase, sialate O-acetylesterase, N-acetylneuraminate lyase, arylesterase, and glycosulfatase activities by strains of fecal bacteria. Infect Immun 60:3971–3978.

71. Deng Z-L, Sztajer H, Jarek M, Bhuju S, Wagner-Döbler I. 2018. Worlds Apart - Transcriptome Profiles of Key Oral Microbes in the Periodontal Pocket Compared to Single Laboratory Culture Reflect Synergistic Interactions. Front Microbiol 9:124.

72. Bubić A, Mrnjavac N, Stuparevic I, Lyczek M, Wielgus-Kutrowska B, Bzowska A, Luic M, Lešcic Ašler I. 2018. In the quest for new targets for pathogen eradication: the adenylosuccinate synthetase from the bacterium Helicobacter pylori. J Enzyme Inhib Med Chem 33:1405–1414.

73. Susin MF, Baldini RL, Gueiros-Filho F, Gomes SL. 2006. GroES/GroEL and DnaK/DnaJ have distinct roles in stress responses and during cell cycle progression in Caulobacter crescentus. J Bacteriol 188:8044–8053.

74. Anglès F, Castanié-Cornet M-P, Slama N, Dinclaux M, Cirinesi A-M, Portais J-C, Létisse F, Genevaux P. 2017. Multilevel interaction of the DnaK/DnaJ(HSP70/HSP40) stress-responsive chaperone machine with the central metabolism. Sci Rep 7:41341.

75. Ogata Y, Mizushima T, Kataoka K, Kita K, Miki T, Sekimizu K. 1996. DnaK heat shock protein of Escherichia coli maintains the negative supercoiling of DNA against thermal stress. J Biol Chem 271:29407–29414.

76. Wong KS, Houry WA. 2012. Novel structural and functional insights into the MoxR family of AAA+ ATPases. J Struct Biol 179:211–221.

77. Coyne MJ, Comstock LE. 2008. Niche-specific features of the intestinal bacteroidales. J Bacteriol 190:736–742.

78. Porter NT, Canales P, Peterson DA, Martens EC. 2017. A Subset of Polysaccharide Capsules in the Human Symbiont Bacteroides thetaiotaomicron Promote Increased Competitive Fitness in the Mouse Gut. Cell Host Microbe 22:494–506.

79. Petit C, Rigg GP, Pazzani C, Smith A, Sieberth V, Stevens M, Boulnois G, Jann K, Roberts IS. 1995. Region 2 of the Escherichia coli K5 capsule gene cluster encoding proteins for the biosynthesis of the K5 polysaccharide. Mol Microbiol 17:611–620.

80. van Selm S, Kolkman MAB, van der Zeijst BAM, Zwaagstra KA, Gaastra W, van Putten JPM. 2002. Organization and characterization of the capsule biosynthesis locus of Streptococcus pneumoniae serotype 9V. Microbiology 148:1747–1755.

81. Dougherty BA, van de Rijn I. 1993. Molecular characterization of hasB from an operon required for hyaluronic acid synthesis in group A streptococci. Demonstration of UDP-glucose dehydrogenase activity. J Biol Chem 268:7118–7124.

82. Cabral DJ, Wurster JI, Belenky P. 2018. Antibiotic Persistence as a Metabolic Adaptation: Stress, Metabolism, the Host, and New Directions. Pharmaceuticals (Basel) 11:14.

83. Rowan AD, Cabral DJ, Belenky P. 2016. Bactericidal antibiotics induce programmed metabolic toxicity. Microb Cell 3:178–180.

84. Allison KR, Brynildsen MP, Collins JJ. 2011. Metabolite-enabled eradication of bacterial persisters by aminoglycosides. Nature 473:216–220.

85. Kohanski MA, Dwyer DJ, Hayete B, Lawrence CA, Collins JJ. 2007. A common mechanism of cellular death induced by bactericidal antibiotics. Cell 130:797–810.

86. Thomas VC, Chittezham Thomas V, Kinkead LC, Janssen A, Schaeffer CR, Woods KM, Lindgren JK, Peaster JM, Chaudhari SS, Sadykov M, Jones J, AbdelGhani SMM, Zimmerman MC, Bayles KW, Somerville GA, Fey PD. 2013. A dysfunctional tricarboxylic acid cycle enhances fitness of Staphylococcus epidermidis during β-lactam stress. mBio 4:e00437.–13–e00437–13.

87. Adolfsen KJ, Brynildsen MP. 2015. Futile cycling increases sensitivity toward oxidative stress in Escherichia coli. Metab Eng 29:26–35.

88. Cho H, Uehara T, Bernhardt TG. 2014. Beta-lactam antibiotics induce a lethal malfunctioning of the bacterial cell wall synthesis machinery. Cell 159:1300–1311.

89. Zinöcker MK, Lindseth IA. 2018. The Western Diet-Microbiome-Host Interaction and Its Role in Metabolic Disease. Nutrients 10:365.

90. Sanchez KK, Chen GY, Schieber AMP, Redford SE, Shokhirev MN, Leblanc M, Lee YM, Ayres JS. 2018. Cooperative Metabolic Adaptations in the Host Can Favor Asymptomatic Infection and Select for Attenuated Virulence in an Enteric Pathogen. Cell 175:146–158.e15.

91. Chassaing B, Van de Wiele T, Gewirtz A. 2017. O-013 Dietary Emulsifiers Directly Impact the Human Gut Microbiota Increasing Its Pro-inflammatory Potential and Ability to Induce Intestinal Inflammation. Inflammatory Bowel Diseases 23.

92. Collins J, Robinson C, Danhof H, Knetsch CW, van Leeuwen HC, Lawley TD, Auchtung JM, Britton RA. 2018. Dietary trehalose enhances virulence of epidemic Clostridium difficile. Nature 553:291–294.

93. Human Microbiome Project Consortium. 2012. A framework for human microbiome research. Nature 486:215–221.

94. Singh RK, Chang H-W, Yan D, Lee KM, Ucmak D, Wong K, Abrouk M, Farahnik B, Nakamura M, Zhu TH, Bhutani T, Liao W. 2017. Influence of diet on the gut microbiome and implications for human health. J Transl Med, 4 ed. 15:73–17.

95. Johnson AJ, Vangay P, Al-Ghalith GA, Hillmann BM, Ward TL, Shields-Cutler RR, Kim AD, Shmagel AK, Syed AN, Personalized Microbiome Class Students, Walter J, Menon R, Koecher K, Knights D. 2019. Daily Sampling Reveals Personalized Diet-Microbiome Associations in Humans. Cell Host Microbe 25:789–802.e5.

96. Turnbaugh PJ, Ley RE, Mahowald MA, Magrini V, Mardis ER, Gordon JI. 2006. An obesity-associated gut microbiome with increased capacity for energy harvest. Nature 444:1027–131.

97. Turnbaugh PJ, Bäckhed F, Fulton L, Gordon JI. 2008. Diet-induced obesity is linked to marked but reversible alterations in the mouse distal gut microbiome. Cell Host Microbe 3:213–223.

98. Thompson LR, Sanders JG, McDonald D, Amir A, Ladau J, Locey KJ, Prill RJ, Tripathi A, Gibbons SM, Ackermann G, Navas-Molina JA, Janssen S, Kopylova E, Vázquez-Baeza Y, Gonzalez A, Morton JT, Mirarab S, Zech Xu Z, Jiang L, Haroon MF, Kanbar J, Zhu Q, Jin Song S, Kosciolek T, Bokulich NA, Lefler J, Brislawn CJ, Humphrey G, Owens SM, Hampton-Marcell J, Berg-Lyons D, McKenzie V, Fierer N, Fuhrman JA, Clauset A, Stevens RL, Shade A, Pollard KS, Goodwin KD, Jansson JK, Gilbert JA, Knight R, Earth Microbiome Project Consortium. 2017. A communal catalogue reveals Earth’s multiscale microbial diversity. Nature 104:11436.

99. Wang Q, Garrity GM, Tiedje JM, Cole JR. 2007. Naive Bayesian classifier for rapid assignment of rRNA sequences into the new bacterial taxonomy. Appl Environ Microbiol 73:5261–5267.

100. McIver LJ, Abu-Ali G, Franzosa EA, Schwager R, Morgan XC, Waldron L, Segata N, Huttenhower C. 2018. bioBakery: a meta’omic analysis environment. Bioinformatics 34:1235–1237.

101. Bolger AM, Lohse M, Usadel B. 2014. Trimmomatic: a flexible trimmer for Illumina sequence data. Bioinformatics 30:2114–2120.

102. Langmead B, Salzberg SL. 2012. Fast gapped-read alignment with Bowtie 2. Nat Methods 9:357–359.

103. Pruesse E, Quast C, Knittel K, Fuchs BM, Ludwig W, Peplies J, Glöckner FO. 2007. SILVA: a comprehensive online resource for quality checked and aligned ribosomal RNA sequence data compatible with ARB. Nucleic Acids Research 35:7188–7196.

104. Wood DE, Salzberg SL. 2014. Kraken: ultrafast metagenomic sequence classification using exact alignments. Genome Biol 15:R46.

105. Lu J, Breitwieser FP, Thielen P, Salzberg SL. 2017. Bracken: estimating species abundance in metagenomics data. PeerJ Computer Science 3:e104.

106. McMurdie PJ, Holmes S. 2013. phyloseq: an R package for reproducible interactive analysis and graphics of microbiome census data. PLoS ONE 8:e61217.

107. Benjamini Y, Hochberg Y. 1995. Controlling the False Discovery Rate: A Practical and Powerful Approach to Multiple Testing. Journal of the Royal Statistical Society Series B (Methodological) 57:289–300.

108. Westreich ST, Korf I, Mills DA, Lemay DG. 2016. SAMSA: a comprehensive metatranscriptome analysis pipeline. BMC Bioinformatics 17:399.

109. Zhang J, Kobert K, Flouri T, Stamatakis A. 2014. PEAR: a fast and accurate Illumina Paired-End reAd mergeR. Bioinformatics 30:614–620.

110. Overbeek R, Olson R, Pusch GD, Olsen GJ, Davis JJ, Disz T, Edwards RA, Gerdes S, Parrello B, Shukla M, Vonstein V, Wattam AR, Xia F, Stevens R. 2014. The SEED and the Rapid Annotation of microbial genomes using Subsystems Technology (RAST). Nucleic Acids Research 42:D206–14.

111. Buchfink B, Xie C, Huson DH. 2015. Fast and sensitive protein alignment using DIAMOND. Nat Methods 12:59–60.

112. Witten IH, Bell TC. 1991. The zero-frequency problem: estimating the probabilities of novel events in adaptive text compression. IEEE Trans Inform Theory 37:1085–1094.

113. Segata N, Izard J, Waldron L, Gevers D, Miropolsky L, Garrett WS, Huttenhower C. 2011. Metagenomic biomarker discovery and explanation. Genome Biol 12:R60.

114. Bushnell B. 2014. BBMap. Genomics of Energy Environment Meeting. Walnut Creek, CA.

115. Li H, Durbin R. 2010. Fast and accurate long-read alignment with Burrows-Wheeler transform. Bioinformatics 26:589–595.

